# The Role of Emotions in Moral Judgments: Time-resolved evidence from event-related brain potentials

**DOI:** 10.1101/541342

**Authors:** Ronja Demel, Michael Waldmann, Annekathrin Schacht

## Abstract

The influence of emotion on moral judgments has become increasingly prominent in recent years. While explicit normative measures are widely used to investigate this relationship, event-related potentials (ERPs) offer the advantage of a preconscious method to visualize the modulation of moral judgments. Based on Gray and Wegner’s (2009) *Dimensional Moral Model*, the present study investigated whether the processing of neutral faces is modulated by moral context information. We hypothesized that neutral faces gain emotional valence when presented in a moral context and thus elicit ERP responses comparable to those established for the processing of emotional faces. Participants (*N*= 26, 13 female) were tested with regard to their implicit (ERPs) and explicit (morality rating) responses to neutral faces, shown in either a morally positive, negative, or neutral context. Higher ERP amplitudes in early (P100, N170) and later (EPN, LPC) processing stages were expected for harmful/helpful scenarios compared to neutral scenarios. Agents and patients were expected to differ for moral compared to neutral scenarios. In the explicit ratings neutral scenarios were expected to differ from moral scenarios. In ERPs, we found indications for an early modulation of moral valence (harmful/helpful) and an interaction of agency and moral valence after 80-120 ms. Later time sequences showed no significant differences. Morally positive and negative scenarios were rated as significantly different from neutral scenarios. Overall, the results indicate that the relationship of emotion and moral judgments can be observed on a preconscious neural level at an early processing stage as well as in explicit judgments.

---

In our daily lives, we are constantly confronted with morally discerning behaviors such as being honest or helping others without personal gain. Morality is not limited to our own actions. It also plays a crucial role in judging others’ behaviors and has been subject to discussion ever since the question of how to lead a righteous life. Rationalist models have long dominated research on moral judgments. They postulated that moral reasoning is a conscious and controlled process. For instance, Piaget (1933) argued that children begin to realize when situations are handled in a manner that seems fair, reasonable or beneficial to all parties by the age of ten and expand their understanding of fairness to include ideal reciprocity (i.e., consideration of another person’s interests; Golden Rule) by middle adolescence. Kohlberg (1969) investigated adolescents’ abstract reasoning abilities. Within these lines, he proposed a new methodology for measuring moral judgment behavior. Through using moral quandaries (e.g., Should Heinz steal a drug to save his wife?), he established a framework for moral reasoning and an assessment tool for moral competencies.

The role of emotions in the moral domain was not only highly neglected over the centuries but has also been the source of large controversies in moral philosophy and psychology. Philosophers, such as Kant, argued that morality is rather based on reason than emotions. Kant’s assumption is based on the *Categorical Imperative* where he states that a rational being could figure out what is the right action to choose by applying the rule “I should never act in such a way that I could not also will that my maximum should be a universal law” (Kant, 1785/1959, p. 18). A violation of the *Categorical Imperative* is declared as immoral. Contradicting Kant’s view, Hume argued that “moral judgments are derived from sentiment, not reason, and we attain moral knowledge by an ‘immediate feeling and finer internal sense,’ not by a ‘chain of argument and induction’” (Hume, 1777/1960, p. 2). He thus assumed a considerable impact of emotion on moral judgments that surpasses the importance of reason. Equally to moral philosophy, most psychological models saw moral judgments as a product of rational reasoning (e.g., Kohlberg, 1969). Haidt (2001) challenged this approach by arguing that moral judgments are generated by intuitive processes. According to his *Social Intuitionist Model (SIM)*, moral intuitions (which include moral emotions) directly cause moral judgments. Decisions are therefore made automatically and a plausible reasoning is only applied *ex post facto*. He proposed this model on the basis of his research which showed that moral judgments are generally best predicted by participants’ affective reactions (Haidt, Koller, & Dias, 1993). Greene et al. (2001) investigated the influence of emotion on moral judgments using functional Magnetic Resonance Imaging (fMRI) and concluded in his *Dual Process Theory* that in moral dilemmas individuals must compromise between the emotion and rationality based cognitive processes that compete in moral reasoning. A first judgment is fast and driven by emotion. Controlled cognitive processes can subsequently change this judgment.. The tight link between morality and emotion that has been proposed in the models was subsequently supported by a large amount of studies which showed that emotion is a key element of moral behavior in humans (Horberg, Oveis, & Keltner, 2011; Tangney, Stuewig, & Mashek, 2007; Waldmann, Nagel, & Wiegmann, 2012).

In line with Haidt’s *SIM* and Greene’s *dual process theory* Gray and Wegner (2011) proposed a specific model of moral emotions that follow the constitution of morality. Moral emotions as a fundamental concept can be divided into a dimensional structure, which is similar to more general models of emotion (e.g., Lang, 1995; Russell, 2003) and are therefore best described by two dimensions: the valence of the emotion that is performed (harm/help) and the moral type (agent/patient). They further argued that a moral action always requires at least two different persons: an agent and a patient. Agency is a position where a person has the capacity to do right or wrong; patiency is the capacity to be a recipient of the wrong or right doing (Gray & Wegner, 2009). Gray and Wegner (2011) assumed four different groups of emotions triggered by four kinds of agents/patients. First, there are villains who elicit anger or disgust and heroes who elicit inspiration and elevation. Second, there are beneficiaries who receive help and therefore induce emotions of relief or happiness and victims who receive a harmful act and are thus related to feelings of sympathy and sadness. Although Gray and Wegner (2011) admitted that any categorical mapping is incomplete, the structure of their model presents a solid basis to investigate the specific emotions elicited by those four kinds of persons described.

Faces are a common source to express and recognize emotions in others in natural environments. Because of their transferability to a scientific setting, they are often used in affective research, both to measure elicitation and recognition of emotion. Even though emotional facial recognition is seen as an automatic and universal process (Ekman, 1992), several studies have shown that the perception of faces highly depends on contextual information, which is particularly relevant in social situations (Barrett, Mesquita, & Gendron, 2011; Carroll & Russell, 1996; Wieser & Brosch, 2012). Davis, Johnstone, Mazzulla, Oler, and Whalen (2010) showed that neutral faces gain affective valence by being paired with emotionally valent context information. In other words, faces previously seen in a positive emotional context evoke a different neural response as compared to those previously seen in either a negative or a neutral context (Morel, Beaucousin, Perrin and George, 2012). Emotion effects are also evident in paradigms were there was no emotional information available through the presented faces but simply through the given non-emotional context information (Schwarz, Wieser, Gerdes, Mühlberger, & Pauli, 2013). Studies have demonstrated that context information is integrated in an unintentional, uncontrollable and effortless manner in face perception (Aviezer, Dudarev, Bentin & Hassin, 2011) and that neutral faces connected to negative context information are classified as more negative compared to neutral faces with neutral context information (Suess, Rabovsky and Rahman, 2014). In a newly published study, Kanunikov and Pavlova (2017) showed differences in event-related potentials (ERPs) for neutral faces that gained valence through a short video clips in which the identities were presented as villains or culprits. Compared to participants who had not seen the clip and also in comparison to other neutral faces, ERPs were significantly amplified for faces that were presented in an emotional context through the video. Given the evidence, it can be assumed that context information influences face perception and is relevant while perceiving inherently neutral faces.

In behavioral studies, we rely on participants’ answers to conclude how a judgment is made and can only assess the end product of a variety of sub processes. Since emotional processing is fast and automatic, it is not reflected in explicit judgments. ERPs can give us much faster and preconscious details compared to explicit judgments and are therefore a fitting method to visualize emotional face processing in dependency of moral context information. To date, only few studies have investigated the relationship of morality and emotion using ERPs (e.g., Cui, Ma, & Luo, 2016; Leuthold, Kunkel, Mackenzie, & Filik, 2014; Luo et al., 2013). Early ERP components are the P100 which was shown to be modulated by facial expressions (e.g. Hammerschmidt, Sennhenn-Reulen, & Schacht, 2017; Rellecke, Palazova, Sommer, & Schacht, 2011) and the N170 that has been linked to structural face encoding (Herzmann, Kunina, Sommer, & Wilhelm, 2010) and as being modulated by emotional expressions (Batty & Taylor, 2003). In later time sequences, there are particular patterns in higher-order processing depending on the specific content of the stimulus (e.g., angry, happy; Hammerschmidt et al., 2017; Schacht & Sommer, 2009). For instance, the P200 and the late positive potential (LPP) are known to be sensitive to emotions. The P200 was shown to be more sensitive to emotionally arousing words when compared to neutral words (Kissler, Assadollahi, & Herbert, 2006), to moral statements clashing with ones own value system (Van Berkum, Holleman, Nieuwland, Otten, & Murre, 2009), and to be related to socio-normative evaluation and conflict detection (Chen, Qiu, Li, & Zhang, 2009; Leuthold et al., 2014). The LPP was shown to be enhanced in response to emotional faces (Schacht & Sommer, 2009) and modulated by preceding narratives (MacNamara, Foti, & Hajcak, 2009).

Integrating the *SIM* (Haidt, 2001) and the *dual process theory* (Greene et al., 2001) our core hypothesis is that the processing of a moral scenario first includes a first appraisal process of valence, followed by a rapid (automatic) emotional response. Subsequently, it comes to higher-order elaborative evaluations of the moral scenarios that finally result in explicit moral judgments. In line with these assumptions, we hypothesize that the ERPs as an implicit measurement differ significantly depending on the valence of the corresponding scenario; a fast, automatic evaluation is expected for the positive and negative scenarios compared to the neutral scenarios. Agents (villains and heroes) in comparison to patients (victims and beneficiaries) are expected to show higher ERPs amplitudes for positive and negative scenarios, whereas for the neutral scenarios, regardless of the person being an agent or a patient, lower ERPs are to be observed. On a behavioral level, we expect that positive and negative scenarios differ significantly from neutral scenarios concerning their morality rating.

## Method

### Participants

Data was collected from 26 healthy participants (13 female, *age range*= 19–30 years, *M_age_=* 23.77, *SD=* 3.15 years). According to self-reports, participants had no neurological or psychiatric disorders and all participants were right-handed (Oldfield, 1971).

### Stimuli and Procedure

The study included face stimuli of the Göttingen Face Database of 399 neutral faces (Kulke, Janßen, Demel, & Schacht, 2017) and 90 moral scenarios that were validated in a different study on the dimension of valence, moral wrongness and plausibility.

The procedure for each trial was as follows: First, a black fixation cross was presented for 1.000ms. Then, two faces were presented simultaneously (one male, one female). Underneath the faces, two names, matching for gender, were randomly inserted from a list of common German names. After a minimum presentation time of 3.000ms and a maximum viewing time of 7.000ms a randomly drawn scenario of an agent and a patient interacting was shown. The names of agent and patient matched the identities that were introduced through picture and corresponding name. The gender of agent and patient was randomized throughout conditions. Each scenario, name, and face stimuli were used only in one trial during the experiment. The allocation of the scenario, the name, and the faces were randomly assigned but matched for gender. To ensure that participants memorized the identities and the scenarios, a minimum viewing time was set to 7.000ms. After a maximum time of 20.000ms, the next screen was displayed automatically. Another fixation cross was presented at the center of the screen for 1.000ms to ensure eye focus on the position of the following target face. The target stimulus was one of the previously introduced identities and was presented centered for 1.000ms. Conditions were randomly presented and equally distributed so that half of the face were female and half of the faces were an agent. After the presentation of the target face, another fixation cross was shown for 1.000ms. Afterwards the question *“How do you judge the behavior from A towards B?”* was shown and had to be answered on a seven-point Likert scale ranging from *1= very harmful* to *7= very helpful.* To ensure that the participants kept paying attention and memorizing the faces that were shown with the scenarios, a one-back face memory task was randomly posed in 20% of the trials over the whole experiment. In the one-back task, participants were shown a face stimulus that was either included in the preceding trial or not included in the whole experiment and were asked the question *“Was this person part of the preceding trial?”*. Their responses were recorded by clicking the *yes* or *no* button below the face stimulus. Feedback about correctness of the answer was given. For each correct answer participants gained an additional bonus of 30 cents. A trial scheme is presented in Figure 2.

**Figure 1.**
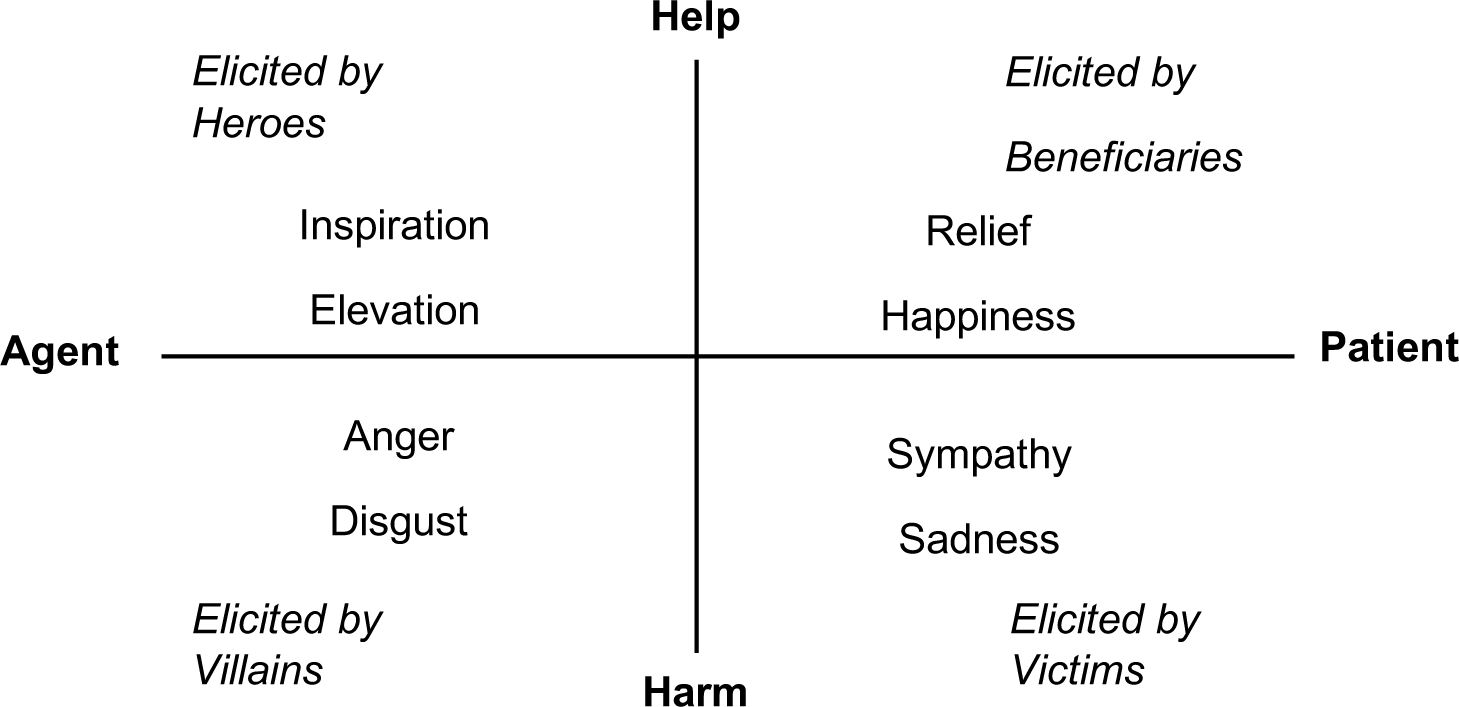
The Dimensional Moral Model (Gray and Wegner, 2011). The structure of moral emotions is mapped by valence (harm/help) and moral type (agent/patient). Emotions in each quadrant are elicited by their respective exemplar.

**Figure 2.**
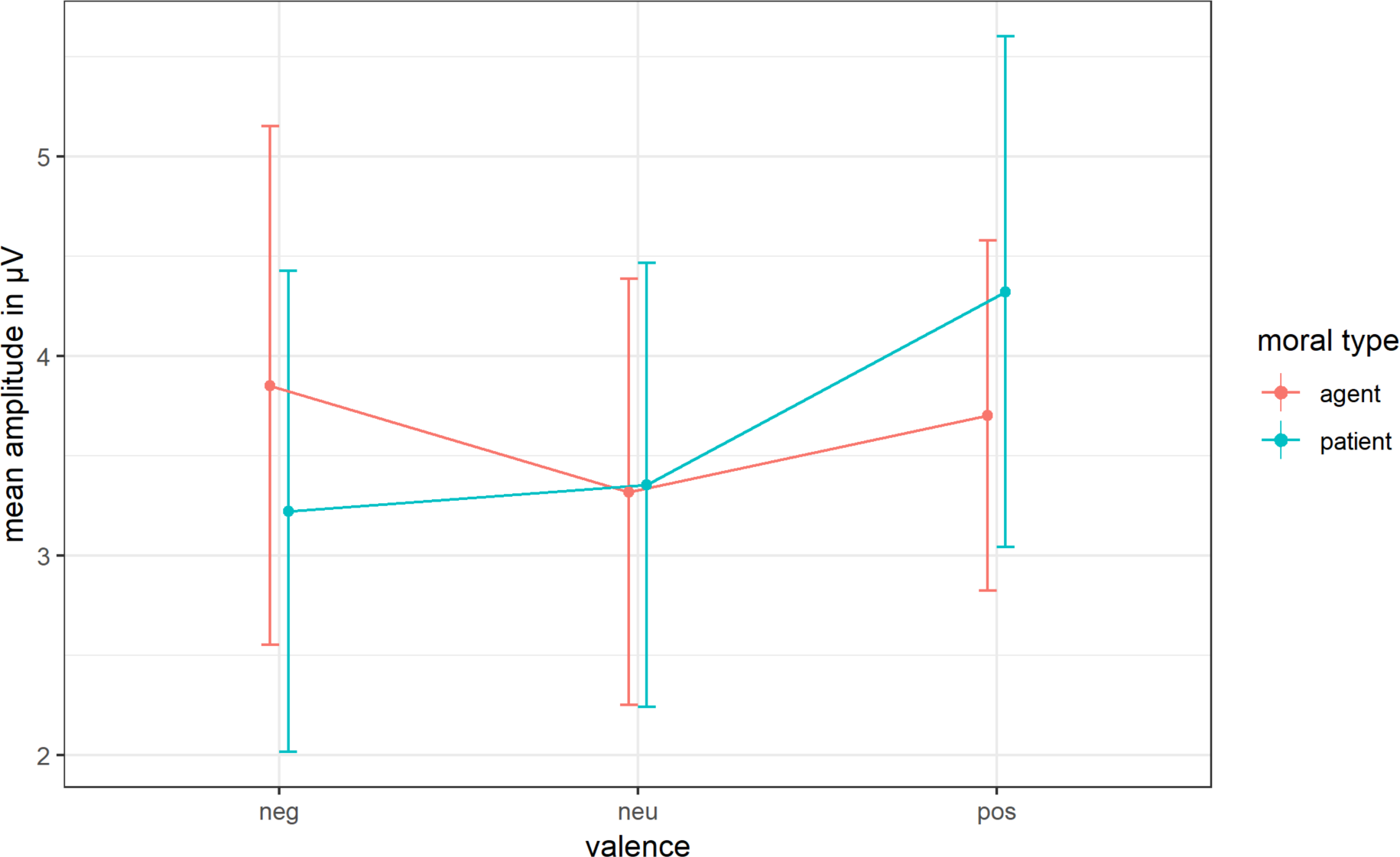
Line chart showing the effects of valence on the mean amplitudes for the P100. On the x-axis the valence categories are displayed. The y-axis shows the mean amplitudes in μV. Effects are depicted for the mean amplitudes and CI for agents and patients. The effects are shown for the time sequence of 80-120 ms.

### Data Recording and Analysis

Python-based visual stimulus system PsychoPy (Peirce, 2007) was used to present and record behavioral data. The EEG was recorded from 64 electrodes mounted in an electrode head cap (Easy-Cap, Biosemi, Amsterdam, Netherlands). The common mode sense (CMS) active electrode and the driven right leg (DRL) passive electrode were used as reference and ground electrodes (cf. www.biosemi.com/faq/cms&drl.htm). Four external electrodes were placed laterally (HEOG) and inferior (VEOG) to the eyes to record eye movements and blinks. Two external electrodes were applied to left and right mastoids (A1, A2). Signals were recorded at a sampling rate of 512 Hz and were amplified with a band pass filter of 0.16-100 Hz. Electrode offsets were kept within a range of ± 20 μV.

Data was processed using Brain Vision Analyzer (Brain Products GmbH, Munich, Germany). The data was re-referenced to average and corrected for blinks using a method of removal of ocular artifact as proposed by Gratton, Coles, & Donchin (1983). The continuous EEG signal was segmented into epochs of 1.200 ms starting 200 ms before stimulus onset and referred to a 100 ms pre-stimulus baseline. After eliminating epochs containing artifacts, segments were averaged per subject and experimental condition (negative agent, negative patient, neutral agent, neutral patient, positive agent, positive patient).

Three early time-windows from 80 to 120 ms (P100 component; Batty & Taylor, 2003), from 140–174 (N170 component; Schacht & Sommer, 2009), and from 200– 250 ms (P200 component; Cui et al., 2016; Leuthold et al., 2014), and a later time-window from 350–600 ms after stimuli onset (LPP component; Cui et al., 2016) were selected for analysis of mean amplitudes. For every investigated ERP component (P100, N170, P200 and LPP) regions of interest (ROIs) were defined through visual inspection and previous evidence. For the P100 and the P200 component, nine electrodes in the occipital-temporal region were selected symmetrically (P7, P8, PO7, PO8, P9, P10, O1, O2, IZ). For the N170 component, ROI was defined using eight temporal-parietal electrodes (P7, P8, P9, P10, PO7, PO8, TP7, TP8). In order to investigate the modulation of the LPP, seven electrodes were used as ROI (CP1, P1, POZ, PZ, CPZ, CP2, P2). The mean amplitudes for each time window were analyzed with LMMs including the fixed factors *valence* (neutral, negative, positive) and *moral type* (agent, patient) and the random factor participant ID.

## Results

### Behavioral Data

Analysis for the morality rating showed means of 1.70 (*SD*= 0.79) for the negative, 4.58 (*SD*= 0.81) for the neutral and 6.28 (*SD*= 0.87) for the positive scenarios from *1= very harmful* to *7= very helpful.* The LMM showed that the valence of the scenarios was significantly different, *F*(2, 2312)= 6334.1*, p<* .001. *η*_*p*_^*2*^= 0.96 The mean scores for negative scenarios were significantly lower when compared to neutral scenarios, *t*(2312)= –69.89, *p<* .001, whereas positive scenarios had significantly higher scores than the neutral scenarios *t*(2312)= 41.46, *p<* .001. All t-tests were Bonferroni corrected.

Error rates in the one-back task were small over all subjects (mean error percentage: 4.92%). This showed that participants looked at the faces attentively and that participants did not have to be excluded due to inattentive execution of the task.

### P1

A LMM was fitted for the time window of 80-120 ms in the parietal P1 ROI. There was no main effect for moral type, *F*(1,125)< 0.00; *p=* .968, *η*_*p*_^*2*^< 0.01. The results for the main effect of valence *F*(1,125)= 2.95, *p=* .055, *η*_*p*_^*2*^=0.01 and the interaction of moral type ✕ valence, *F*(1,125)= 2.40, *p=* .094, *η*_*p*_^*2*^=0.01 were also not significant, although they indicated a trend. *Figure 3* depicts the mean amplitudes and confidence intervals (CI) for agents and patients in each valence category. Planned pairwise comparisons of the means are reported in Table 2.

**Table 1.** Examples of the scenarios.

|  |  |
| --- | --- |
| Help | Anna hits a tree with her car and the car catches on fire immediately. Peter, who is driving towards the accident scene, stops and pulls Anna out of the car before it explodes.<br>(Anna fährt mit dem Auto gegen einen Baum. Das Auto beginnt sofort zu brennen. Peter fährt auf den Unfall zu, hält an und zieht Anna aus dem brennenden Auto, bevor das Auto explodiert.) |
| Neutral | Lena and Nico attend the same literature club ever since their first semester. Nico received a book from his grandmother for Christmas. Lena asks him if he liked it.<br>(Lena und Nico besuchen seit ihrem ersten Semester gemeinsam einen Literaturclub. Nico hat zu Weihnachten ein Buch von seiner Oma geschenkt bekommen. Lena fragt ihn, ob es ihm gefallen hat.) |
| Harm | Hanna asks Anton to take a letter (containing important application documents) to the post office for her. Anton throws the letter in the trash instead of taking it to the post office.<br>(Hanna bittet Anton einen Brief für ihn einzuwerfen, der wichtige Bewerbungsunterlagen enthält. Anton schmeißt den Brief in den Müll, anstatt ihn zur Post zu bringen.) |

**Table 2.** z-values and p-values of the planned pairwise comparisons of the means for the P100 component.

| condition 1 | condition 2 | z | p |
| --- | --- | --- | --- |
| negative | neutral | 0.72 | .473 |
|  | positive | -1.73 | .084 |
| neutral | positive | -2.44 | .014 * |
| agent | patient | 0.15 | .882 |
| negative agent | neutral agent | 1.12 | .264 |
| negative patient | neutral patient | -0.10 | .918 |
| negative agent | positive agent | 0.45 | .652 |
| negative patient | positive patient | -2.90 | .004 ** |
| neutral agent | positive agent | -0.67 | .505 |
| neutral patient | positive patient | -2.79 | .005 ** |

**Figure 3.**
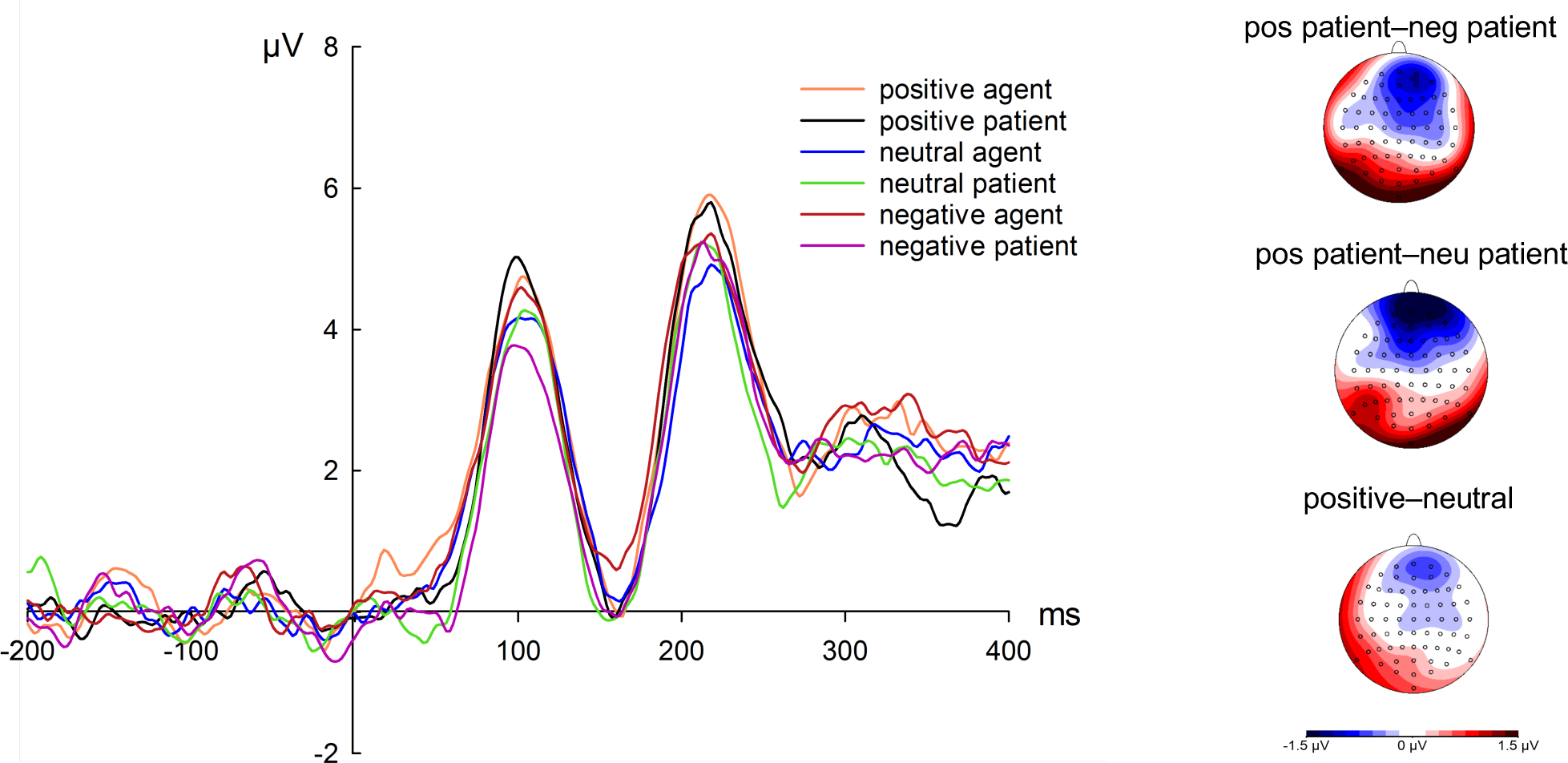
Grand-average ERP waveforms at O1/O2 electrode site and ERP difference maps for significant comparisons relative to the target face onsets.. The voltage scale ranges from – 1.5 μV to 1.5 μV

### N170, P200, LPP

The analysis using LMMs for the respective ROI and time sequences of the N170 (140–174 ms), the P200 (200–250 ms), and the LPP (350–600 ms) revealed no significant main effects or interactions for the valence and the moral type (see Table 3).

**Table 3.** Main effects and interactions for the N170, P200 and LPP components.

| Components | | *df* | *F* | *p* | $\eta_p^2$ |
| --- | --- | --- | --- | --- | --- |
| *N170* | Valence | 125 | 0.53 | .587 | <0.01 |
|  | Moral type | 125 | 2.12 | .147 | <0.01 |
|  | Valence × Moral type | 125 | 0.04 | .952 | <0.01 |
| *P200* | Valence | 125 | 1.76 | .175 | <0.01 |
|  | Moral type | 125 | 0.01 | .909 | <0.01 |
|  | Valence × Moral type | 125 | 0.20 | .818 | <0.01 |
| *LPP* | Valence | 125 | 0.29 | .748 | <0.01 |
|  | Moral type | 125 | 1.74 | .188 | 0.01 |
|  | Valence × Moral type | 125 | 1.55 | .215 | 0.01 |

*Note:* *df*, *F*-value, *p*-value and partial $\eta^2$ are presented for the main effect of valence, moral type and the interaction effect valence x moral type. None of the presented comparisons reached significance.

## Discussion

The present study aimed at investigating the connection of morality and emotion using the underlying structure of a model that has been introduced by Gray and Wegner (2011). The first hypothesis postulated a difference in the three valence categories introduced through the scenarios, which resulted in differences in the ratings and a difference within the ERP components. Results showed that the three valence categories were understood in terms of morality. The actions of villains were rated as significantly more harmful than the actions of neutral agents, whereas the actions of positive agents were rated as significantly more helpful than the actions of neutral agents. Concerning the ERP components, there was a trend for valence on the early time sequence of 80–120 ms after stimulus onset. This points towards a fast, emotional evaluation of morally laden faces depending on the valence category.

The second hypothesis concerned the influence of moral type (agent/patient) when evaluating the target face. It was expected that agents elicit larger amplitudes in ERPs compared to patients. Since there were no significant effects for the later ERP components only assumptions based on a trend in the P100 component can be made. Planned contrasts showed that there is no difference between agents and patients. Therefore, the hypothesis cannot be confirmed.

Although valence was only induced implicitly the observed trend for valence indicates an emotional modulation in the early time sequence from 80 to 120 ms after stimulus onset. Planned comparisons of the means showed that positively associated identities elicit larger mean amplitudes compared to neutrally associated identities. This is in line with the hypothesis that valent scenarios elicit higher mean amplitudes than neutral scenarios, however this effect could not be found for the comparison of the negatively with the neutrally connoted faces. Rozin and Royzman (2001) have argued that negative information is weighed more strongly than positive information in moral judgments which in consequence would suggest a preferential and more pronounced processing of morally negative connoted faces. In contrast, Zhao and collegues (2017) found a positivity bias for face processing depending on the manipulation of direct threat, and high vs. low ability context information. Thus, context information seems to have an influential impact on the processing of morally valent information. There also was a trend for the interaction of moral type ✕ valence. Planned contrasts of means showed that positive patients differed significantly from neutral and negative patients. The higher mean amplitudes for positive patients compared to neutral and negative patients further emphasize the preferential processing of a face that is associated with being a beneficiary. To explain this outcome, it is assumed that for positive patients (beneficiaries) and negative agents (villains) the expected behavior is clearer, whereas, for positive agents (heroes) and negative patients (victims), the expected reaction is ambiguous. Villains can be labeled easily as being morally reprehensible and beneficiaries as deserving. This might result in less conflict and, therefore, in higher mean amplitudes. As for heroes and victims, this relationship is more complex. The heroes’ act implies a general sense of “ought”, meaning it is an obligation to act in a helping way when being confronted with a situation where one can prevent another person being harmed. Supererogatory acts, as in doing something that is beyond expectation, are rare in real life. In the set of scenarios presented in the present experiments, there is also an underrepresentation of superogatory acts. For victims, it is possible that one assumes reasons why the person is deserving of not being helped, which has also been described in the theory of *moral disengagement* (Bandura, 2002).

The trend for valence for of the P100 component indicated a very early emotional evaluation of the target face. The effect of valence in early ERP components such as the P100 point to a rapid emotional modulation (e.g., Aguado et al., 2012; Rossi et al., 2017; Schacht & Sommer, 2009; Stolarova, Keil, & Moratti, 2006) allowing the fast salience detection of the most relevant stimuli for the preparation of an adequate response (Hammerschmidt et al., 2017). Our study supports this line of reasoning. Emotional content was processed extremely fast even though it was only associated with the face via the coupled scenario. From an evolutionary perspective, deciding appropriately on emotionally salient stimuli is an essential tool to survival. It is however remarkable that this ability seems to be extended to stimuli that are only emotionally relevant through their context information.

For the analysis of the N170, the P200 and the LPP components, three time windows were selected. A negativity after 140–174 ms was observed for all conditions and the scalp distribution of this sequence matched the topography of other studies (e.g., Morel et al., 2012; Wieser & Brosch, 2012). There also was a positive peak after 200 ms. For the LPP, the scalp distribution showed similarities to previous studies that investigated emotional processing (e.g., Cui et al., 2016; Schacht & Sommer, 2009). However, for these ERP components (N170, P200, and the LPP), no significant main effects or interactions were found. This contradicts our hypotheses. It was expected that there is an emotional modulation of the ERP components– especially for the LPP since this component has been associated with higher-order processing. Other studies also reported the absence of later emotionally related ERP components after finding an early neural modulation of faces (P100; Aguado et al., 2012; Hammerschmidt et al., 2017). It remains open whether the null effects represent a non-difference of the valence categories or, rather, if the association of face and corresponding scenario was not processed as an identity in higher order mechanisms. In other words, the association of the face with the background information might not have been strong enough to cause a higher order processing of morality.

Overall, in our experimental design the cognitive load that participants had to process while executing the task was high. Working memory load has been reported to alter ERP components in different processing stages (e.g., N170, EPN, LPP; Lin, Schulz, & Straube, 2016; Morgan, Klein, Boehm, Shapiro, & Linden, 2008; Van Dillen & Derks, 2012). For instance, Lin, Schulz, and Straube (2016) found that congruency effects in ERP responses on emotional faces are modulated by cognitive load during the expectation phase. Effects of cognitive load are also shown in early components such as the N170. Therefore, it is expected that visual areas play an important role for the working memory regarding faces (Morgan et al., 2008). Similar effects of high cognitive load might have altered the effects in the present ERP study. Participants had to uphold the name, facial information and information of two identities through the scenarios per trial, which might have demanded a high load of working memory capacity during the task.

The present studies used a model by Gray and Wegner (2011). The authors recognized that the categorical mapping of their model is abstract and therefore lacking in some parts. This results in good and bad characters without any gray scaling. The scenarios that we used to illustrate an identity for the faces only depict the identity in one situation. Since characters are more complex and not situation-dependent, one scenario or one moral act might have provided only very little information to form a valid impression. This link might have been too weak to elicit higher ERP effects. Apart from the very brief description of one situation, a lot of imagination was demanded from the participants. They had to integrate the faces, names and background information in a very abstract and artificial manner. One might therefore argue that ecological validity was lacking in the present studies since no actual interaction was shown. There has been evidence that faces are processed differently and also show differences in ERP amplitudes when their naturalness has been altered (Risko, Laidlaw, Freeth, Foulsham, & Kingstone, 2012). Depicting the moral context information might help to form a more natural impression and therefore enhance ecological validity.

## Conclusion

The present study extend the knowledge on the influence of emotion in moral judgments using explicit and implicit measurements. The explicit judgments point to a good understanding for the emotional valence of the moral scenarios. In the ERP study, the modulation of valence and moral type was found at an early processing stage (80–120 ms after stimulus onset). Differences in later time sequences were not significant. Thus, it remains open if there are no effects at higher processing stages or if the used design was not suitable for measuring emotion effects due to its complexity. Emotion effects in moral judgments using ERP technique should be further explored with a simpler design in subsequent studies. This pilot study scrutinized the importance to further investigate the mechanisms of morality in affective face processing.

## References

Aguado, L., Valdés-Conroy, B., Rodríguez, S., Román, F. J., Diéguez-Risco, T., & Fernández-Cahill, M. (2012). Modulation of early perceptual processing by emotional expression and acquired valence of faces: An ERP study. Journal of Psychophysiology, 26(1), 29–41. https://doi.org/10.1027/0269-8803/a000065

Aviezer, H., Dudarev, V., Bentin, S., & Hassin, R. R. (2011). The Automaticity of Emotional Face-Context Integration. Emotion, 11(6), 1406–1414. https://doi.org/10.1037/a0023578.The

Bandura, A. (2002). Selective Moral Disengagement in the Exercise of Moral Agency. Journal of Moral Education, 31(2), 101–119. https://doi.org/10.1080/0305724022014322

Barrett, L. F., Mesquita, B., & Gendron, M. (2011). Context in Emotion Perception. Current Directions in Psychological Science, 20(5), 286–290. https://doi.org/10.1177/0963721411422522

Batty, M., & Taylor, M. J. (2003). Early processing of the six basic facial emotional expressions. Cognitive Brain Research, 17(3), 613–620. https://doi.org/10.1016/S0926-6410(03)00174-5

Carroll, J. M., & Russell, J. A. (1996). Do facial expressions signal specific emotions? Judging emotion from the face in context. Journal of Personality and Social Psychology, 70(2), 205–218. https://doi.org/10.1037/0022-3514.70.2.205

Chen, P., Qiu, J., Li, H., & Zhang, Q. (2009). Spatiotemporal cortical activation underlying dilemma decision-making: An event-related potential study. Biological Psychology, 82(2), 111–115. https://doi.org/10.1016/j.biopsycho.2009.06.007

Cui, F., Ma, N., & Luo, Y.-J. (2016). Moral judgment modulates neural responses to the perception of other’s pain: an ERP study. Scientific Reports, 6, 20851. https://doi.org/10.1038/srep20851

Davis, F. C., Johnstone, T., Mazzulla, E. C., Oler, J. A., & Whalen, P. J. (2010). Regional response differences across the human amygdaloid complex during social conditioning. Cerebral Cortex, 20(3), 612–621. https://doi.org/10.1093/cercor/bhp126

Ekman, P. (1992). An argument for basic emotions. Cognition & Emotion. https://doi.org/10.1080/02699939208411068

Gratton, G., Coles, M. G. H., & Donchin, E. (1983). A new method for off-line removal of ocular artifact. Electroencephalography and Clinical Neurophysiology, 55(4), 468–484. https://doi.org/10.1016/0013-4694(83)90135-9

Gray, K., & Wegner, D. M. (2009). Moral typecasting: divergent perceptions of moral agents and moral patients. Journal of Personality and Social Psychology, 96(3), 505–520. https://doi.org/10.1037/a0013748

Gray, K., & Wegner, D. M. (2011). Dimension of Moral Emotions. Emotion Review, 3(3), 258–260. https://doi.org/10.1177/1754073911402388

Greene, J. D., Sommerville, R. B., Nystrom, L. E., Darley, J. M., & Cohen, J. D. (2001). An fMRI Investigation of Emotional Engagement in Moral Judgment. Science, 293(5537), 2105–2108. https://doi.org/10.1126/science.1062872

Haidt, J. (2001). The Emotional Dog and Its Rational Tail: A Social Intuitionist Approach to Moral Judgment. Psychological Review, 108(4), 814–834. https://doi.org/10.1037//0033-295X.

Haidt, J., Koller, S. H., & Dias, M. G. (1993). Affect, culture, and morality, or is it wrong to eat your dog? Journal of Personality and Social Psychology, 65(4), 613–628. https://doi.org/10.1037/0022-3514.65.4.613

Hammerschmidt, W., Sennhenn-Reulen, H., & Schacht, A. (2017). Associated Motivational Salience Impacts Early Sensory Processing of Human Faces. NeuroImage, (April), 0–1. https://doi.org/10.1016/j.neuropsychologia.2017.01.017

Herzmann, G., Kunina, O., Sommer, W., & Wilhelm, O. (2010). Individual differences in face cognition: brain-behavior relationships. Journal of Cognitive Neuroscience, 22(3), 571–589. https://doi.org/10.1162/jocn.2009.21249

Horberg, E. J., Oveis, C., & Keltner, D. (2011). Emotions as Moral Amplifiers: An Appraisal Tendency Approach to the Influences of Distinct Emotions upon Moral Judgment. Emotion Review, 3(3), 237–244. https://doi.org/10.1177/1754073911402384

Hume, D. (1777). An enquiry concerning the principles of morals. La Salle, IL, US: Open Court.

Kant, I. (1785). Foundation of the metaphysics of morals. (L. Beck, Ed.). Indianopolis, IN, US: Bobbs-Merrill.

Kanunikov, I. E., & Pavlova, V. I. (2017). Event-Related Potentials to Faces Presented in an Emotional Context. Neuroscience and Behavioral Physiology, 967–975. https://doi.org/10.1007/s11055-017-0498-8

Kissler, J., Assadollahi, R., & Herbert, C. (2006). Emotional and semantic networks in visual word processing: insights from ERP studies. Progress in Brain Research, 156(06), 147–183. https://doi.org/10.1016/S0079-6123(06)56008-X

Kohlberg, L. (1969). Stage and sequence: The cognitive-developmental approach to socialization. In D. Goslin (Ed.), Handbook of socialization theory and research (pp. 347–480).

Kulke, L., Janßen, L., Demel, R., & Schacht, A. (2017). Validating the Goettingen Faces Database. Open Science Framework. https://doi.org/http://doi.org/10.17605/OSF.IO/4KNPF

Lang, P. J. (1995). The Emotion Probe: Studies of Motivation and Attention. American Psychologist, 50(5), 372–385. https://doi.org/10.1037/0003-066X.50.5.372

Leuthold, H., Kunkel, A., Mackenzie, I. G., & Filik, R. (2014). Online processing of moral transgressions: ERP evidence for spontaneous evaluation. Social Cognitive and Affective Neuroscience, 10(8), 1021–1029. https://doi.org/10.1093/scan/nsu151

Lin, H., Schulz, C., & Straube, T. (2016). Effects of expectation congruency on event-related potentials (ERPs) to facial expressions depend on cognitive load during the expectation phase. Biological Psychology, 120(2016), 126–136. https://doi.org/10.1016/j.biopsycho.2016.09.006

Luo, Y., Shen, W., Zhang, Y., Feng, T. yong, Huang, H., & Li, H. (2013). Core disgust and moral disgust are related to distinct spatiotemporal patterns of neural processing: An event-related potential study. Biological Psychology, 94(2), 242–248. https://doi.org/10.1016/j.biopsycho.2013.06.005

MacNamara, A., Foti, D., & Hajcak, G. (2009). Tell me about it: neural activity elicited by emotional pictures and preceding descriptions. Emotion, 9(4), 531–543. https://doi.org/10.1037/a0016251

Morel, S., Beaucousin, V., Perrin, M., & George, N. (2012). Very early modulation of brain responses to neutral faces by a single prior association with an emotional context: Evidence from MEG. NeuroImage, 61(4), 1461–1470. https://doi.org/10.1016/j.neuroimage.2012.04.016

Morgan, H. M., Klein, C., Boehm, S. G., Shapiro, K. L., & Linden, D. E. J. (2008). Working memory load for faces modulates P300, N170, and N250r. Journal of Cognitive Neuroscience, 20(6), 989–1002. https://doi.org/10.1162/jocn.2008.20072

Oldfield, R. C. (1971). The assessment and analysis of handedness: The Edinburgh inventory. Neuropsychologia, 9(1), 97–113. https://doi.org/10.1016/0028-3932(71)90067-4

Peirce, J. W. (2007). PsychoPy-Psychophysics software in Python. Journal of Neuroscience Methods, 162(1–2), 8–13. https://doi.org/10.1016/j.jneumeth.2006.11.017

Piaget, J. (1933). The Moral Judgment of the Child. Journal of Educational Psychology, 24(2), 157–158. https://doi.org/10.1037/h0067118

Rellecke, J., Palazova, M., Sommer, W., & Schacht, A. (2011). On the automaticity of emotion processing in words and faces: Event-related brain potentials evidence from a superficial task. Brain and Cognition, 77(1), 23–32. https://doi.org/10.1016/j.bandc.2011.07.001

Risko, E. F., Laidlaw, K., Freeth, M., Foulsham, T., & Kingstone, A. (2012). Social attention with real versus reel stimuli: toward an empirical approach to concerns about ecological validity. Frontiers in Human Neuroscience, 6(May), 1–11. https://doi.org/10.3389/fnhum.2012.00143

Rossi, V., Vanlessen, N., Bayer, M., Grass, A., Pourtois, G., & Schacht, A. (2017). Motivational Salience Modulates Early Visual Cortex Responses across Task Sets. Journal of Cognitive Neuroscience, 26(6), 968–979. https://doi.org/10.1162/jocn

Rozin, P., & Royzman, E. B. (2001). Negativity Bias, Negativity Dominance, and Contagion. Personality and Social Psychology Review, 5(4), 296–320. https://doi.org/10.1207/S15327957PSPR0504_2

Russell, J. (2003). Core affect and the psychoogical construction of emotion. Psychological Review, 110, 145–172.

Schacht, A., & Sommer, W. (2009). Emotions in word and face processing: Early and late cortical responses. Brain and Cognition, 69(3), 538–550. https://doi.org/10.1016/j.bandc.2008.11.005

Schwarz, K. A., Wieser, M. J., Gerdes, A. B. M., Mühlberger, A., & Pauli, P. (2013). Why are you looking like that? How the context influences evaluation and processing of human faces. Social Cognitive and Affective Neuroscience, 8(4), 438–445. https://doi.org/10.1093/scan/nss013

Stolarova, M., Keil, A., & Moratti, S. (2006). Modulation of the C1 visual event-related component by conditioned stimuli: Evidence for sensory plasticity in early affective perception. Cerebral Cortex, 16(6), 876–887. https://doi.org/10.1093/cercor/bhj031

Tangney, J. P., Stuewig, J., & Mashek, D. J. (2007). Moral Emotions and Moral Behavior. Annual Reviews, 58, 345–372. https://doi.org/10.1146/annurev.psych.56.091103.070145.Moral

Van Berkum, J. J. A., Holleman, B., Nieuwland, M., Otten, M., & Murre, J. (2009). Right or Wrong? The Brain’s Fast Response to Morally Objectionable Statements. Psychological Science, 20(9), 1092–1099. https://doi.org/10.1111/j.1467-9280.2009.02411.x

Van Dillen, L. F., & Derks, B. (2012). Working memory load reduces facilitated processing of threatening faces: An ERP study. Emotion, 12(6), 1340–1349. https://doi.org/10.1037/a0028624

Waldmann, M. R., Nagel, J., & Wiegmann, A. (2012). Moral Judgment BT - The Oxford Handbook of Thinking and Reasoning. The Oxford Handbook of Thinking and Reasoning, (19), 274–299. https://doi.org/10.1093/oxfordhb/9780199734689.001.0001

Wieser, M. J., & Brosch, T. (2012). Faces in context: A review and systematization of contextual influences on affective face processing. Frontiers in Psychology, 3(NOV), 1–13. https://doi.org/10.3389/fpsyg.2012.00471

Zhao, S., Xiang, Y., Xie, J., Ye, Y., Li, T., & Mo, L. (2017). The Positivity Bias Phenomenon in Face Perception Given Different Information on Ability. Frontiers in Psychology, 8(April), 1–8. https://doi.org/10.3389/fpsyg.2017.00570

